# Cell cycle independent role of cyclin D3 in host restriction of SARS-CoV-2 infection

**DOI:** 10.1101/2022.05.07.491022

**Authors:** Ravi K. Gupta, Petra Mlcochova

**Affiliations:** Cambridge Institute of Therapeutic Immunology & Infectious Disease (CITIID), Cambridge, UK; Department of Medicine, University of Cambridge, Cambridge, UK; Africa Health Research Institute, Durban, South Africa

**Author notes:** **Material and Correspondence** should be addresses to Petra Mlcochova,.

## Abstract

The COVID-19 pandemic caused by Severe Acute Respiratory Syndrome Coronavirus 2 (SARS-CoV-2) presents a great threat to human health. The interplay between the virus and host plays a crucial role in successful virus replication and transmission. Understanding host-virus interactions is essential for development of new COVID-19 treatment strategies. Here we show that SARS-CoV-2 infection triggers redistribution of cyclin D1 and cyclin D3 from the nucleus to the cytoplasm, followed by its proteasomal degradation. No changes to other cyclins or cyclin dependent kinases were observed. Further, cyclin D depletion was independent from SARS-CoV-2 mediated cell cycle arrest in early S phase or S/G2/M phase. Cyclin D3 knockdown by small interfering RNA specifically enhanced progeny virus titres in supernatants. Finally, cyclin D3 co-immunoprecipitated with SARS-CoV-2 Envelope and Membrane proteins. We propose that cyclin D3 inhibits virion assembly and is depleted during SARS-CoV-2 infection to restore efficient assembly and release of newly produced virions.

## Introduction

The severe acute respiratory syndrome coronavirus 2 (SARS-CoV-2) is the causative agent for the global Covid-19 pandemic. To date, SARS-CoV-2 has infected over 265 millions of people with death toll of more than 5 million people ^1^. While strategies of counteracting SARS-CoV-2 infection through vaccination have been partially successful, there is still a need for effective antiviral drugs given the emergence of vaccine escape variants such as Omicron. Coronaviruses, including SARS-CoV-2, like other viruses are intracellular pathogens exploiting the host cell machinery to their own advantage. The identification of cellular mechanisms and host cell targets required for SARS-CoV-2 life cycle will provide us with new knowledge that could be used to interfere with viral replication and therefore presents an alternative approach to block viral infection.

Cyclins and cyclin dependent kinases (CDKs) are the major regulators of cell cycle progression. Many viruses, including coronaviruses, adopt a strategy of manipulating cell cycle progression through cyclin-CDKs complexes ^2-5^ to facilitate viral replication. Several SARS-CoV-1 proteins have been shown to reduce cyclin D and cyclin E and A expression that is connected to cell cycle arrest ^6-8^. For example, SARS-CoV-1 N protein directly interacts with cyclin D to prolong the S phase ^6^ that ensure enough supply of nucleotides for viral replication. Nsp13 protein both in SARS-CoV-1 and Infectious bronchitis virus (IBV) interacts with DNA polymerase subunit to induce DNA damage and cell cycle arrest ^9^. It is believed that the virus infection associated cell cycle arrest increases essential DNA repair processes and replication proteins that are required by virus replication.

A recent study showed that SARS-CoV-2 infection is correlated with cell arrest at S/G2 transition based on a comparison of phosphoproteomic profiles of SARS-CoV-2 infected VERO E6 cells and phosphorylation profiles collected at specific cell cycle phases. Further by measuring DNA content an increase in the fraction of cells in S and G2/M phase with decreased proportion of cells in G0/G1 phase has been observed. Additionally, kinase activity profiling uncovered that CDK1/2 activities are reduced by SARS-CoV-2, possibly adding to S/G2 phase arrest ^10^.

Here we present additional comprehensive data on SARS-CoV-2 cell cycle changes and identified cyclin D3 as a novel interactor with SARS-CoV-2 viral proteins. We propose that SARS-CoV-2 effectively reduces cyclin D3 levels in infected cells to achieve efficient viral assembly.

## Results

### SARS-CoV-2 infection depletes cyclin D1 and D3

Several coronaviruses are known to regulate cell cycle and cell cycle associated proteins, including cyclins ^2,3,5,7,8,11^. To determine whether the reduction in cyclins can occur during productive SARS-CoV-2 infection, the abundance of cyclins and cell cycle associated proteins (Fig. 1) from SARS-CoV-2 infected cells were compared to uninfected VERO AT2 cells (Fig. 1B-D) and human epithelial cell line A549 AT2 (Fig. 1E,F). Western blots (Fig. 1B, E) and densitometric analysis (Fig. S1A,B) indicate that cyclin D1 and D3 levels are significantly reduced compared to uninfected cells or cells infected with heat inactivated virus. Several other cyclins tested did not show any changes in expression nor did cell cycle associated kinases that form a functional complex with cyclins and regulate together cell cycle (Fig.1A,B).

**Figure 1.**
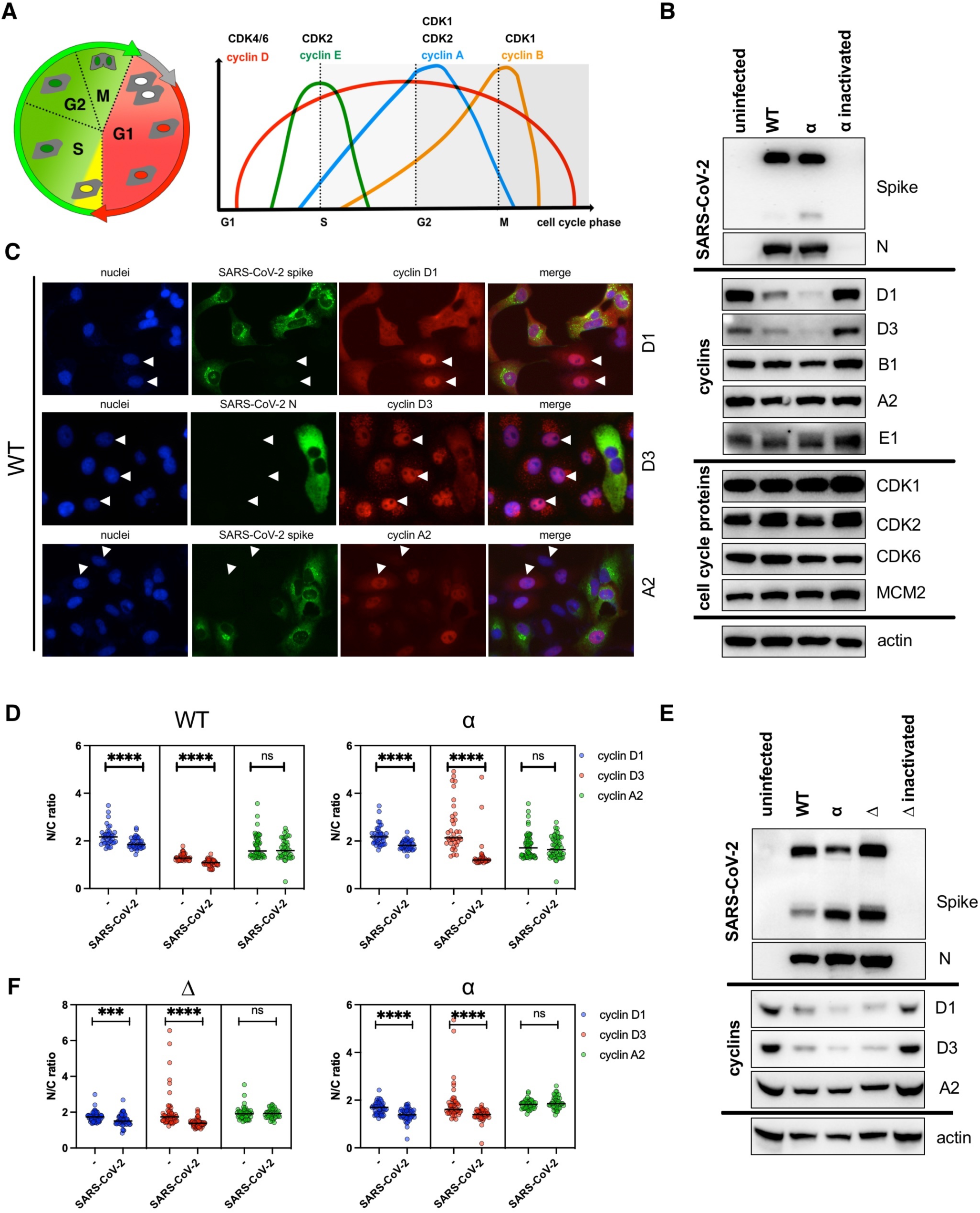
SARS-CoV-2 infection depletes cyclin D1 and D3. (A) Diagram of cell cycle and cyclin expression during cell cycle. Cyclins drive cell cycle changes by interacting with cyclin dependent kinases (CDK). (B-C) VERO AT2 cells were infected with WT and alpha (α) SARS-CoV-2 live virus variants, and heat inactivated α variant at MOI 0.1. (B) Cells were lysed 48h post-infection and viral protein as well as cell cycle associated protein expression was analysed by western blot. N, nucleocapsid. (C) VERO AT2 Cells were fixed 24h post-infection and stained for viral proteins and cyclins. Arrowheads highlight un-infected cells and cyclin D/A nuclear localization. Arrowheads: Nuclear cyclin staining in uninfected cells. (D) VERO AT2 cells. Quantification of cyclin A2, D1 and D3 relocalization from nucleus after infection. Uninfected (-) and SARS-CoV-2 infected cells were identified by negative/positive nucleocapsid or Spike staining. Ratio between nuclear and cytoplasm staining intensity of cyclins was measured using ImageJ and Harmony (PerkinElmer). At least 50 cells were counted. Statistical analysis was performed using two-sided unpaired Student’s t-tests; ns, non-significant; ****p < 0.0001. (E) A549 AT2 cells were infected with WT, alpha (α), and delta (Δ) SARS-CoV-2 variants. Cells were lysed 48h post-infection and viral, cyclin proteins expression was analysed by western blot. N, nucleocapsid. (F) A549 AT2 cells. Quantification of cyclin A2, D1 and D3 relocalization from nucleus after infection. Uninfected (-) and SARS-CoV-2 infected cells were identified by negative/positive nucleocapsid or Spike staining. Ratio between nuclear and cytoplasm (N/C ratio) staining intensity of cyclins was measured using ImageJ and Harmony (PerkinElmer). At least 50 cells were counted. Statistical analysis was performed using two-sided unpaired Student’s t-tests; ns, non-significant; ***p < 0.001; ****p < 0.0001.

We examined the cellular distribution of cyclins D1, D3 and A2 in infected VERO AT2 cells by immunofluorescence analysis (Fig.1C, S1C). The ratio between the fluorescence intensity of cyclins in the nucleus and cytoplasm (N/C ratio) indicates that while uninfected cells localise cyclin D1/D3 predominantly in nucleus (higher ratios), these cyclins are relocalised to the cytoplasm in infected cells (lower ratios) (Fig. 1C,D,F). Importantly, cyclin A2 that was not degraded in infected cells (Fig. 1B), and did not differ in cellular localisation between infected and uninfected cells. Relocation of cyclin D3 was also confirmed in HeLa cells expressing ACE2 (Fig. S2).

These data indicate that a productive SARS-CoV-2 infection leads to both reduced levels of cyclin D1 and D3 and their cellular relocalisation.

### Proteasome inhibition abolishes effect of SARS-CoV-2 infection on D-cyclins depletion

It is known that D-cyclins are degraded mainly through the ubiquitin dependent 26S proteasomal degradation pathway ^12,13^. To further investigate the mechanism of D-cyclins depletion during SARS-CoV-2 infection, we investigated the involvement of proteasome degradation pathway. Cells were infected in the absence or presence of protease inhibitors MG-132 and Bortezomib and D-cyclin levels were subsequently measured (Fig. 2, S3). Firstly, the addition of both inhibitors before or early after infection blocked SARS-CoV-2 infection (Fig. S3A). This is in concordance with the published inhibitory effect of MG-132 on SARS-CoV-2 Mpro ^14^. In view of these results, A459 AT2 cells were infected with Delta SARS-CoV-2 variant for 24h, before addition of the proteasome inhibitor Bortezomib for an additional 24h. Uninfected cells treated with the same drug showed an increase in D-cyclin levels, suggestive of cyclin stabilization in cells. This increase was more pronounced in the case of cyclin D1 than in D3 which did not reach statistically significant levels (Fig. 2A,B). Importantly, in the absence of proteasome inhibition D-cyclins relocalised from nucleus to cytoplasm and were degraded in infected cells. But proteasome inhibition significantly stabilized D-cyclins levels in infected cells where D-cyclins expression levels were equal to those in uninfected cells (Fig 2A,B,S3B,C). Further, proteasome inhibition also prevented relocalisation of cyclin D3 to the cytoplasm in SARS-CoV-2 infected VERO AT2 cell (S3B,C). Even though the proteasome inhibition did not completely prevent the translocation of cyclin D3 in infected A549 AT2 cells, it revealed significant presence of cyclin D3 in the nucleus (Fig. 2C,D).

**Figure 2.**
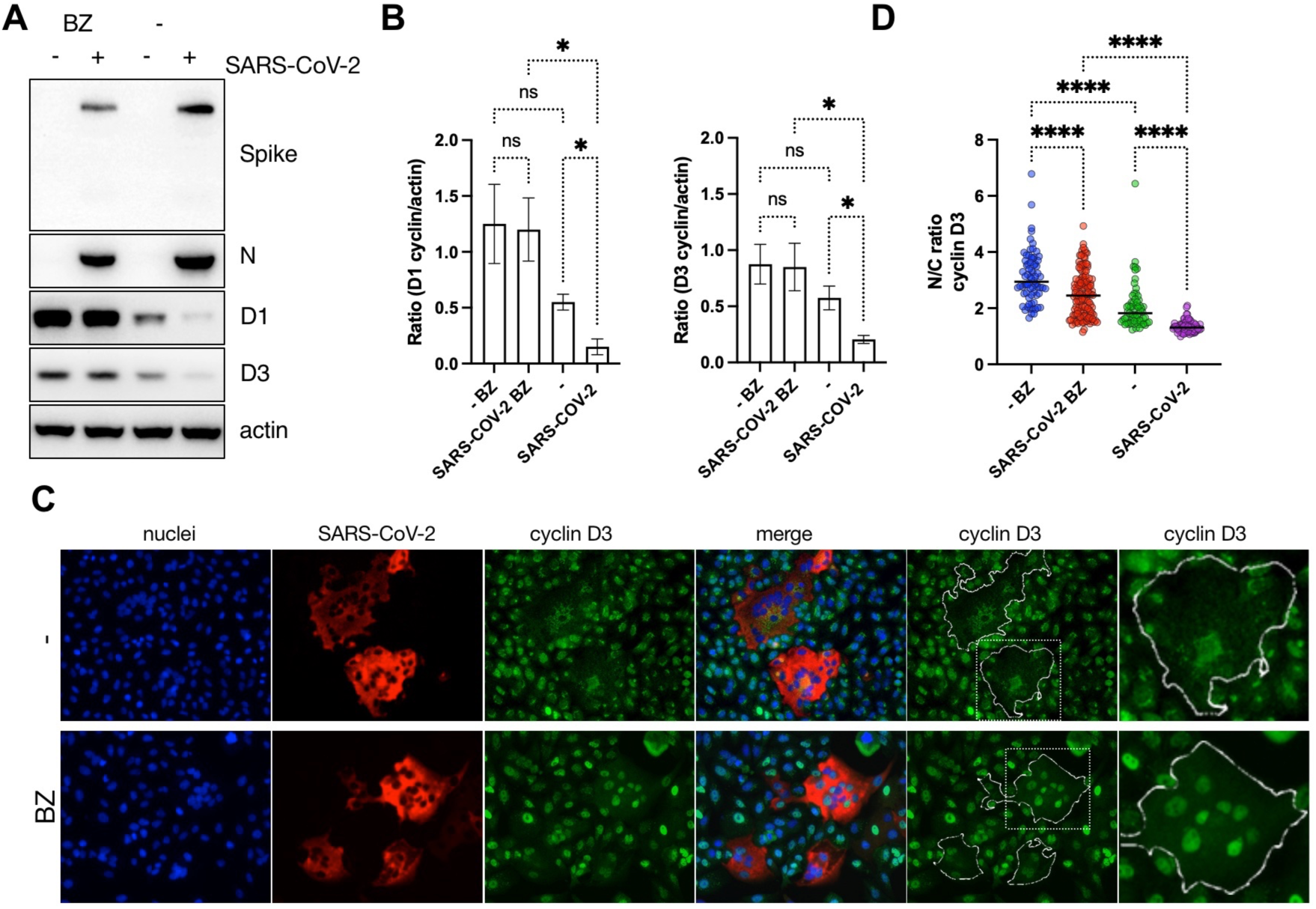
Proteasome inhibition abolishes effect of SARS-CoV-2 infection on D-cyclins depletion. (A) A549 AT2 cells were infected with Delta SARS-CoV-2 variant. Proteasome inhibitor Bortezomib (BZ, 1uM) was added to cells 18h post-infection. Cells were lysed 24h post-addition of inhibitor and cyclins and viral proteins were detected by western blot. (N) nucleocapsid. (B) Densitometry analysis of western blots for D-cyclins (normalized to actin) in A549 AT2 cells. Plots are average of 3 independent experiments. Bars indicate mean with SD. Statistical analysis was performed using ordinary two way ANOVA; ns, non-significant; *p < 0.1. (C) A459 AT2 cells were infected with Delta SARS-CoV-2 variant. Proteasome inhibitor Bortezomib (BZ, 1uM) was added to cells 8h post-infection. Cells were fixed and stained 24h post-addition of inhibitor. (D) Quantification of D3 cyclin relocalization from nucleus after infection. Uninfected (-) and SARS-CoV-2 infected cells were identified by negative/positive nucleocapsid staining. Ratio between nuclear and cytoplasm (N/C ratio) staining intensity of cyclins was measured using ImageJ and Harmony (PerkinElmer). At least 50 cells have been counted. Statistical analysis was performed using ordinary two-way ANOVA; ***p < 0.001.

These data indicate that the D-cyclin depletion during SARS-CoV-2 infection is mediated a proteasome-dependent pathway.

### Cyclin D3 negatively regulates SARS-CoV-2 infection

To understand the functional role of D-cyclins in SARS-CoV-2 pathogenesis, the effect of cyclin knockdown on viral replication was investigated. siRNA effectively reduced levels of D and A2 cyclins by >80% at 48h post-transfection, compared to non-targeting control (NT) in A549 AT2 cells (Fig.3A,S4A,B) or VERO AT2 (Fig.3C). Cells depleted for individual cyclins were infected with Delta, Alpha and WT variant to allow multiple rounds of infection (Fig 3, S4). Interestingly, only viral titres from cyclin D3-depleted cells were significantly higher than those from control (NT). This was evident for all SARS-CoV-2 variants tested in both A549 AT2 and VERO AT2 cells (Fig. 3B,D). These data support the notion that cyclin D3 negatively modulates SARS-CoV-2 infection and could potentially have impact on viral spread.

**Figure 3.**
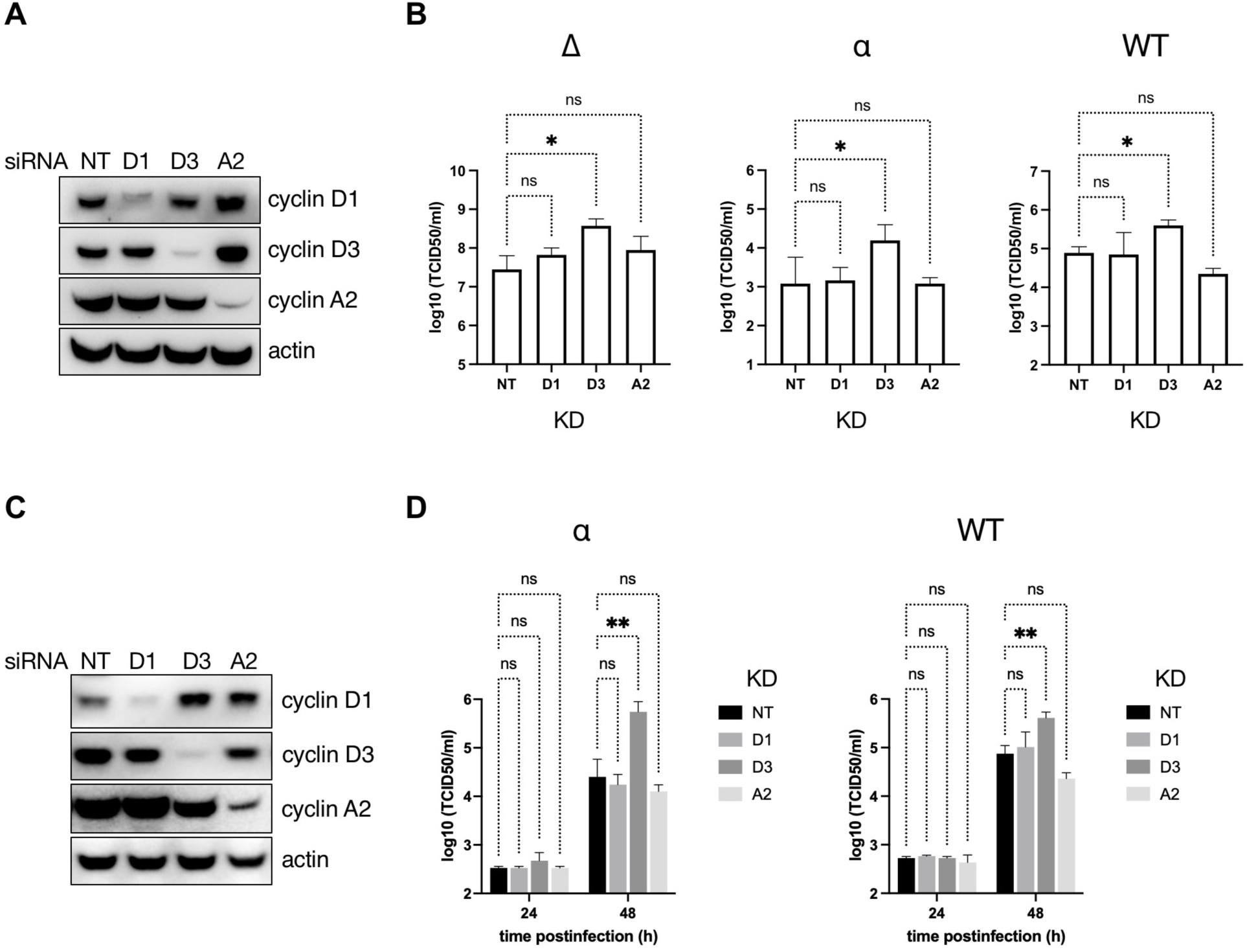
Cyclin D3 impairs SARS-CoV-2 spreading infection. (A,B) D and A-cyclins were depleted using siRNA in A549 AT2 cells. Cells were infected 18h later with Delta (Δ), Alpha (α) or wild type (WT) SARS-CoV-2 variants at MOI 0.001, 0.1, 0.1 respectively. Cells were washed 4h post-infection and new media were added. Supernatants and cells were collected 48h later for western blots (A) and TCID50 (B). (C,D) D and A-cyclins were depleted using siRNA in VERO AT2 cells. Cells were infected 18h later with Alpha (α) or wild type (WT) SARS-CoV-2 variants at MOI 0.1. Cells were washed 4h post-infection and new media added. Supernatants and cells were collected at 24h and 48h later. (A,C) Representative example of western blot from cell lysates collected at 48h post-infection. (B,D) Virus titres in cell culture supernatants were determined as TCID50 in VERO AT2 cells. Graphs represent average of n=3. Statistical analysis was performed using one-way ANOVA with Dunnett’s multiple comparisons test. NT, non-targeting control. ns, non-significant; *p < 0.1; **p < 0.01. Bars indicate mean with SD.

### Productive SARS-CoV-2 infection induces cell cycle arrest

Cyclin D1, D2, and D3 are important regulators of G1 to S phase progression. Given the observed depletion of cyclin D in SARS-CoV-2 infected cells we speculated that this depletion might affect cell cycle progression. Indeed, SARS-CoV-2-mediated S/G2 cell cycle arrest has been reported recently in non-human VERO E6 cells ^10^ but its role in human cells is unknown. To understand the role between infection and D-cyclin depletion in cell cycle regulation we firstly aimed to examine previously reported cell cycle arrest phenotype. We used the FUCCI (fluorescence ubiquitination cell cycle indicator) Cell Cycle Sensor (Fig. S5) ^15^ in VERO AT2 and A549 AT2 cells. VSV-G pseudotyped viral particles containing FUCCI sensor were used to transduce cells. Cells were then infected with a replication competent strain of SARS-CoV-2 for 24h, fixed and stained for nucleocapsid. Flow cytometry analysis was used to visualise G1, early S and S/G2/M phase. The FUCCI system cannot demonstrate G0 phase as it is defined as a cell population void of any fluorescent protein but cannot be differentiated from an untransduced cell population. Cells were gated on infected (expressing SARS-CoV-2 nucleocapsid) or uninfected cells with Cdt1/Geminin expression determined in both gated populations and compared (Fig. 4A-D). Indeed, SARS-CoV-2 infection mediated cycle arrest in S/G2/M phase in VERO AT2 cells (Fig. 4C), confirming previously published data ^10^. Interestingly, SARS-CoV-2 mediated cell cycle arrest in A549 AT2 cells was identified specifically in early S phase (Fig. 4D). This cell cycle arrest was caused by productive SARS-CoV-2 infection as use of heat inactivated virus or Remdesivir (RDV) treatment abrogated cell cycle arrest (Fig. S6A-C) and population of arrested cells increased with increasing MOI (Fig. S6 D,E). All SARS-CoV-2 variants tested (WT, Alpha, Delta) caused cell cycle arrest in S/G2/M phase in VERO AT2 cells and in early S phase in A549 AT2 and (Fig. 4E,F). Furthermore, cell cycle kinetics in virus exposed but uninfected cells were similar to unexposed uninfected cells (Fig. S6 F,G).

**Figure 4.**
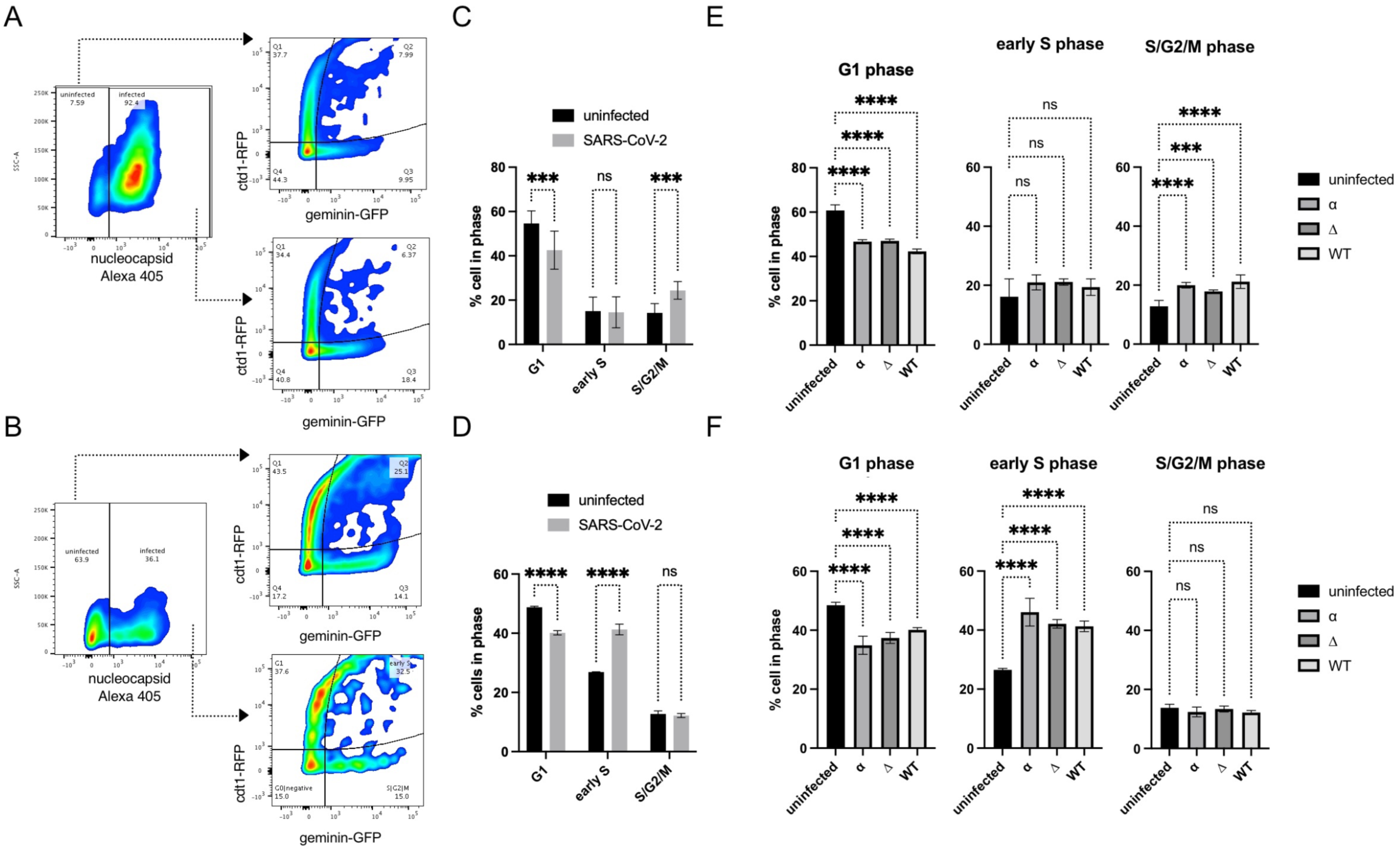
SARS-CoV-2 infection arrests cell cycle. Cells were transduced with Fucci containing lentiviral particles for 18h and infected with SARS-CoV-2 variants for additional 24h. (A) VERO AT2 or (B) A549 AT2 cells. Example of gating strategy for cell cycle analysis. The population of cells exposed to SARS-CoV-2 was stained for nucleocapsid (N) protein and gated on N+ve and N-ve population. Cells were further analysed for expression of cdt1 and Geminin. See also Supplemental Figure S5. (C) VERO AT2 or (D) A549 AT2 cells infected with WT SARS-CoV-2. Quantification of Cdt1 +ve cells (G1), Cdt1/Geminin +ve cells (early S phase) and Geminin +ve cells (S,G2,M phase) cells. n = 5; one-way ANOVA with Dunnett’s multiple comparisons test: ns, non-significant; ****p < 0.0001. Bars indicate mean with SD. (E) VERO AT2 or (F) A549 AT2 cells. Quantification of cell cycle arrest after exposure of cells to SARS-CoV-2 variants. α, alpha (MOI 0.5); Δ, delta (MOI 0.1); WT, Wuhan (MOI 0.5). n = 4; one-way ANOVA with Dunnett’s multiple comparisons test: ns, non-significant; ****p < 0.0001; ***p < 0.001. Bars indicate mean with SD.

These observations support the hypothesis that a productive SARS-CoV-2 infection is responsible for cell cycle arrest and that the arrest is not the result of by-stander effects. However, the data suggest that the specific phase in which cells are arrested appears to be cell-type dependent.

### SARS-CoV-2 mediated Cyclin D depletion is cell cycle independent

To dissect whether the increase in viral titer is a direct consequence of cylin D depletion or aftermath of cell cycle arrest caused by the absence of this cyclin we investigated cell cycle arrest during SARS-CoV-2 infection and linked it to cyclin D expression/cellular localization. It has been previously reported that the Cyclin D degradation is sufficient to cause cell cycle arrest in G1 phase ^16,17^. We confirmed cell cycle arrest in G1 phase in A549 AT2 after D-cyclin depletion (Fig. S7A,B). Further VERO AT2 cells showed similar G1 arrest after individual D1 and D3 cyclin depletion but as well when both D1 and D3 cyclins were depleted together (Fig.S7C). Importantly, no increase in early S nor S/G2/M phase have been observed after cyclin D depletion in uninfected cells. On the contrary, a decrease in these phases has been identified in concordance with more cells arresting in G1 phase and not progressing through the cell cycle (Fig.S7). As we have already shown that SARS-CoV-2 infection increases percentage of cells in early S (in A549 AT2 cells) and S/G2/M (in VERO AT2) specific cell cycle phases (Fig.4), we investigated cell cycle progression in infected cells were D-cyclins and cyclin A had been depleted (Fig. 5, S8). A549 AT2 cells have been depleted for D and A cyclins (Fig. 5A) and infected with Delta variant. The percentage of infected cells was determined 24h later and a small increase (3 fold) compare to NT control was detected in cells depleted for cyclin D3 (Fig.5B,C). The A549 AT2 cell population in early S increased without any changes in S/G2/M phase following infection in both non-targeted (NT) control cells and cells depleted for D and A cyclins (Fig.5D). This data were confirmed using an Alpha SARS-CoV-2 variant (Fig. S8A,B). Furthermore, cell cycle in virus exposed but uninfected cells showed similar cell cycle kinetics as unexposed and uninfected cells excluding any by-stander effect of infection on cell cycle changes (Fig. S8C,D). Moreover, VERO AT2 cells showed an increase in the proportion of cells in S/G2/M phase after SARS-CoV-2 infection even when D and A cyclins were depleted, confirming our results in A549 AT2 cells (Fig. S8E,F). Additionally, single cell analysis showed that D-cyclins are relocated and degraded in SARS-CoV-2 infected cells independently from cell cycle phase. VERO AT2 cells were transduced with Fucci containing lentiviral particles and 18h later infected with Delta (Fig. 5E-G) or Alpha (Fig. S8 E,G,H) SARS-CoV-2 variant. Quantification of nuclear/cytoplasm ratio of D-cyclin staining in different cells cycle phases (identified by expression of Fucci sensor) clearly showed that cyclin D3 (Fig.5F) and cyclin D1 (Fig.5G) in SARS-CoV-2 infected cells are relocalized from nucleus to cytoplasm probably for degradation, independent of the cell cycle phase they are in.

**Figure 5.**
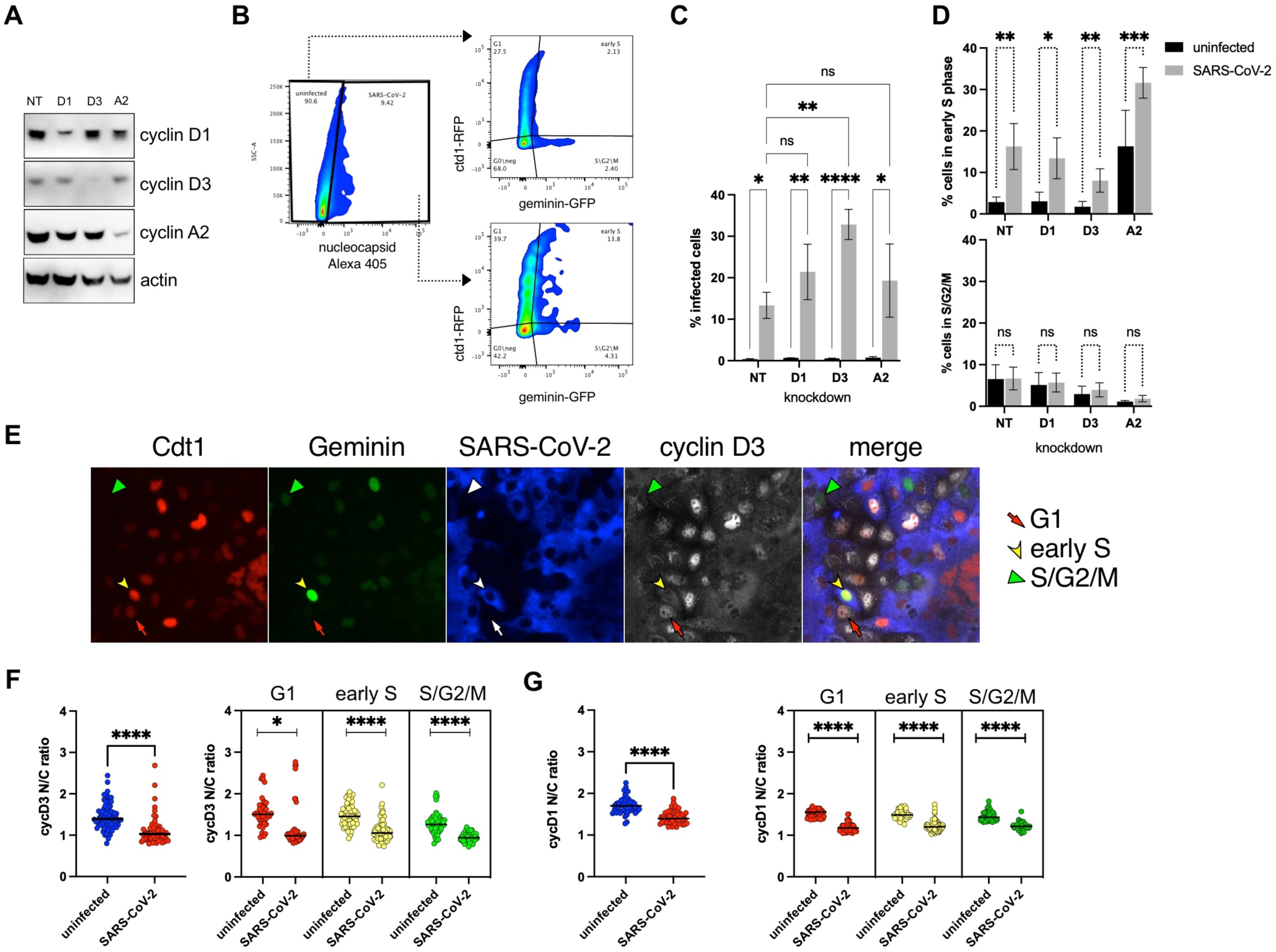
SARS-CoV-2 mediated depletion of D-cyclins is cell cycle arrest independent. (A-D) A549 AT2 cells were depleted for D and A2 cyclins and 18h later infected with Delta variant SARS-CoV-2 for 24h. Cells were fixed, stained for SARS-CoV-2 nucleocapsid and analysed for infection and Fucci cell cycle sensor. (A) A representative western blot from lysates of uninfected knock-down cells. (B) Example of gating strategy for flow cytometry analysis. (C) Percentage of infected cells in cells depleted for cyclins. n=3; Ordinary two-way ANOVA with Sidak’s multiple comparisons test: ns, non-significant; ****p < 0.0001; ***p < 0.001; **p < 0.01; *p < 0.1. Bars indicate mean with SD. (D) Flow cytometry analysis of early S and S/G2/M cell cycle phases comparing cyclin D1, D3, and A2 knockdown to NT (non-target siRNA). n = 3. Statistical analysis was performed using two-sided unpaired Student’s t-tests; ns, non-significant; ***p < 0.001; **p < 0.01; *p < 0.1. Bars indicate mean with SD. (E-G) VERO AT2 cells were transduced with VSV-G pseudotyped Fucci containing lentiviral particles and 18h later infected with Delta variant SARS-CoV-2. Cells were fixed and stained for D-cyclins 24h later. (E) Example of acquisition using automated microscopic platform. Cells are identified for infection, expression of cyclin D3, and cell cycle (Red/arrow=G1phase; Green/arrowhead=S/G2/M; Red+Green/arrowhead=early S). (F-G) Quantification of D-cyclins re-localization from nucleus to cytoplasm and correlation with cell cycle phases using ImageJ and Harmony (PerkinElmer). (F) Cyclin D3. (G) Cyclin D1. At least 50-200 cells were analysed in each condition. Statistical analysis was performed using two-sided unpaired Student’s t-tests; ***p < 0.001; *p < 0.1.

These data suggest that SARS-CoV-2 mediated D-cyclin depletion is cell cycle independent as (i) SARS-CoV-2 infection caused VERO AT2 and A549 AT2 cell arrest in S/G2/M or early S phases, respectively, while D-cyclins were degraded in both cell lines, (ii) depletion of D-cyclins did not cause cells cycle arrest in early S or S/G2/M phase; (iii) based on single cell analysis of Fucci sensor and cyclin D expression and localization, SARS-CoV-2 infection caused D-cyclin nuclear depletion independently from cell cycle phase the cells were in.

### Cyclin D3 associates with E and M SARS-CoV-2 proteins

Cyclin D3 has been previously implicated in the restriction of influenza A virus through impairment of virus assembly ^18^. Cyclin D3 has been shown to interact with IAV protein M2, a ion channel that promotes viral replication ^18,19^. Interestingly, SARS-CoV-2 E protein has been suggested to be an ion channel ^20,21^. In the light of our data showing cyclin D3 depletion increased SARS-CoV-2 viral titer, we investigated potential implication of cyclin D3 in SARS-CoV-2 assembly using immunoprecipitation. Firstly, the interaction between cyclin D3 and E protein was investigated (Fig. 6A). HA-tagged cyclin D3 was co-expressed together with Strep-tagged E and nsp9 protein in 293T cells. Nsp9 was chosen as a control on the basis of its diverse cellular localization both in the nucleus and cytoplasm ^22^. Of note, 293T cells showed undetectable endogenous expression of cyclin D3 (Fig. S9A). We showed that SARS-CoV-2 Envelope (E) coimmunoprecipitated with HA-tagged cyclin D3 using anti-HA antibody while SARS-CoV-2 nsp9 protein did not (Fig. 6A) suggestive of specific binding to E protein.

**Figure 6.**
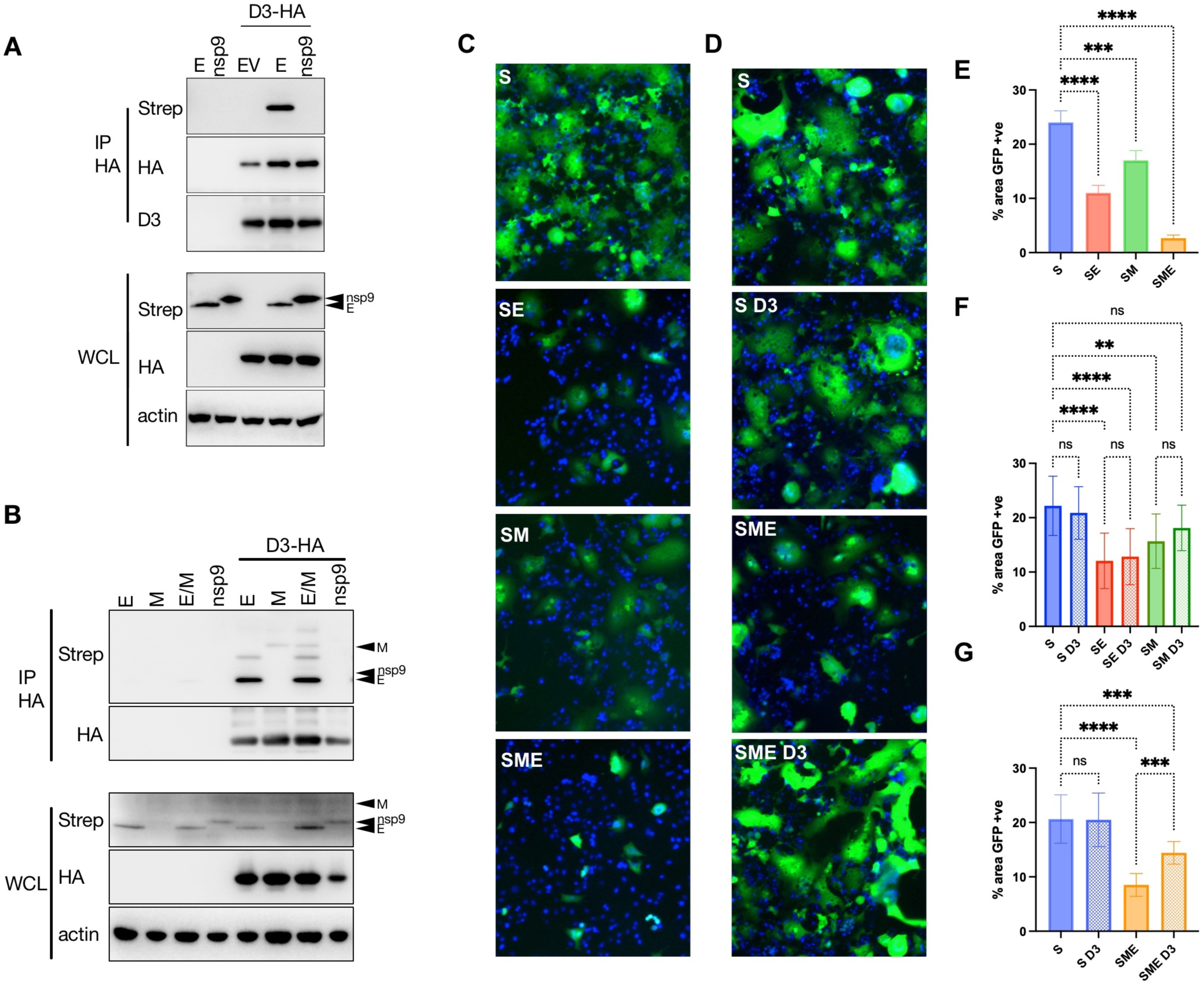
Cyclin D3 associates with SARS-CoV-2 protein E and M. (A) 293T cells were contransfected with HA - cyclin D3 and Strep-tag-SARS-CoV-2 E or nsp9, and control plasmid (EV). Immunoprecipitation was performed using anti-HA antibody. The immunoprecipitates were blotted with anti-Strep, anti-HA, and cyclin D3 antibodies. (B) 293T cells were contransfected with HA - cyclin D3 and Strep-tag-SARS-CoV-2 E, M, both E and M or nsp9. Immunoprecipitation was performed using anti-HA antibody. The immunoprecipitates were blotted with anti-Strep, anti-HA antibodies. WCL, whole cell lysate. (C-G) 293T GFP11 cells were transfected with Spike, and/or with Envelope, Membrane, and cyclin D3. 24h post-transfection cells were seeded at a 1:1 ratio with Vero-GFP10 cells and percentage of GFP+ve area (syncytia) were determined 18h later. S, Spike; E, Envelope; M, Membrane; D3, cyclin D3. (C,D) Representative images of GFP+ syncytia. (E-G) Quantification of cell-to-cell fusion showing percentage of the GFP+ve area to the acquired total cell area. n = 5; one-way ANOVA with Dunnett’s multiple comparisons test: ns, non-significant; ****p < 0.0001; ***p < 0.001; **p < 0.01. Bars indicate mean with SD.

SARS-CoV-2 E protein is a known interactor of Membrane (M) and Nucleocapsid (N) proteins. Further, it has been shown for SARS-CoV-1 and SARS-CoV-2 that membrane (M), Envelope (E) and Nucleocapsid (N) structural proteins are required for efficient assembly ^23-26^. We expressed HA-tagged cyclin D3 with Spike (S), or Strep-tagged N, M, E, nsp9 (Fig. S9). Coimmunoprecipitation with HA-tagged cyclin D3 using anti-HA antibody or anti-cyclin D3 mouse monoclonal antibody revealed an interaction with M and E proteins (Fig. S9 C,D). Further, coimmunoprecipitation using beads binding Strep-tagged protein showed pull down of HA-tagged cyclin D3 with M and E proteins (Fig. S9E). There was no evidence for binding between cyclin D3 and N or S proteins (Fig. S9). Furthermore, Cyclin D3 interaction with M and E was confirmed by expressing cyclin D3 with E or M or both E or M proteins together (E/M) and immunoprecipitated using anti-HA antibody (Fig. 6B).

To further confirm interaction between cyclin D3 and E and M we assessed effect of cyclin D3 expression on E and M function during virus assembly. It has been previously reported that SARS-CoV-2 E and M proteins regulate intracellular trafficking and processing of Spike, leading to S is retention in the ER and Golgi, preventing syncytia formation ^27^. It is possible that this interaction between structural proteins and spike retention allows S to target the virion assembly sites. We hypothesised that if cyclin D3 is interacting with E and M it might impact their function in Spike processing/trafficking, and we can use it as a read-out by assessing the syncytia formation. Firstly, a split GFP system ^28^ was used to confirm that E and M or combination of both (E/M) impact Spike-mediated syncytia formation. Indeed, both structural proteins when coexpressed with S significantly decreased GFP+ve area (cell-cell fusion) (Fig.6C,E). Secondly, 293T GFP11 cells were transfected with full length S (WT), and/or with E, M, and cyclin D3. 24h post-transfection cells were seeded at a 1:1 ratio with Vero-GFP10 and cell to cell fusion was measured 18h later to determine a proportion of green area to total phase area (Fig.6D-G). Interestingly, cyclin D3 had no effect on reduction of syncytia when expressed together with S or S + E or S + M (Fig.6F). However, it increased syncytia formation in combination with S+M+E suggestive of compromising E and M impact on Spike processing/trafficking towards the cell surface (Fig.6D,G).

These data together demonstrate that cyclin D3 associates with proteins important for SARS-CoV-2 assembly and impairs their optimal function.

## Discussion

Here we show that SARS-CoV-2 infection depletes levels of cyclin D and suggest that this depletion is independent from changes to cell cycle arrest in infected cells. Further, cyclin D3 seems to interfere with efficient SARS-CoV-2 assembly by interacting with Envelope and Membrane SARS-CoV-2 proteins.

The molecular mechanism of coronavirus-mediated regulation of cell cycle and cell cycle associated proteins has not been comprehensively investigated, especially in SARS-CoV-2 infection.

Many viruses can manipulate cell cycle of infected cells. Coronaviruses are not an exception ^2-8^. SARS-CoV-2 has been reported to arrest cell cycle in S/G2/M phase in VERO E6 cells ^10^. Our own data confirmed this observation using the Fucci system ^29^, and comparing two different cell lines VERO AT2 (monkey, *Cercopithecus aethiops*, epithelial/kidney) and A549 AT2 (human, epithelial/lung) both expressing ACE2/TMPRSS2, we uncovered that cell cycle arrest in SARS-CoV-2 infected cells occurs at different stages of cell cycle phases. While cell cycle arrest in VERO AT2 is at late S and G2/M phase as previously reported ^10^, A549 AT2 cell are arrested specifically at early S phase. This difference would not be possible to uncover if classical cell cycle techniques, like propidium iodine or DAPI staining, would be used as they can not separate G0 and G1 phase nor early S phase. Based on these data we can conclude that SARS-CoV-2 infection of human cells arrests the cell cycle in S phase, possibly to create a favourable environment for viral replication and spread. However, the exact mechanism is unclear at the present.

Cyclin dependent kinases (CDKs) and corresponding cyclins are essential part of cell cycle progression. Several coronaviruses have been shown to regulate these proteins ^2-8^. Specifically down-regulation of cyclin D1 has been shown in IBV ^5^, and SARS-CoV-1 ^6^, cyclin D3 reduction in SARS-CoV-1 infection ^7^. However, decrease in cyclin Ds has been always connected to cell cycle arrest. Here we show that SARS-CoV-2 mediate translocation of cyclin D1 and D3 from nucleus into cytoplasm for proteasome degradation in both VERO AT2 and A549 AT2 cells. Analysis of other cyclins in VERO AT2 cell did not reveal any changes to cyclin A (promoting S phase entry and mitosis), B (accumulates in G2 phase), or E (limiting factor for G1 phase progression and S phase entry), supporting the notion that degradation of cyclin D proteins is specific.

Further, we present data supporting a notion that the down-regulation of cyclin D during SARS-CoV-2 infection is cell cycle arrest independent. Firstly, detailed analysis of cell cycle arrest in cells depleted for cyclin D1 and D3 revealed arrest in G1 phase as published previously ^16^ not in S or G2/M phase. Nevertheless, when cells in the absence of cyclin Ds were infected, cell were still arrested in early S phase (in A549 AT2) or S/G2/M (in VERO AT2), suggestive of cell cycle arrest during infection being caused by a factor other than depletion of cyclin D proteins. Secondly, using single cell microscopy and Fucci cell cycle sensor allowed us to measure cyclin Ds re-localization from nucleus into cytoplasm for degradation in specific cell cycle phases. All cell cycle phases detected showed that in the presence of infection cyclin Ds are always re-localized from nucleus to cytoplasm for degradation. It has been published recently that CDK1/2 activities are reduced during SARS-CoV-2 infection and might be leading to S/G2 phase arrest ^10^. Although no changes at expression level were evident for CDKs in our study, phosphorylation state of these kinases was not investigated, and it is thus possible that such changes may occur and effect cell cycle progression.

If cyclin D is not used as a mean for virus to arrest cell cycle, it is possible that it might represent a host restriction factor preventing optimal viral replication and spread. Importantly, depletion of cyclin D3 by siRNA increased viral titres after SARS-CoV-2 infection in viral supernatants. This suggest that cyclin D3 might play role in viral spread. This effect is reminiscent of role of cyclin D3 in influenza infection, where cyclin D3 depletion resulted in increase viral production. This study also showed that cyclin D3 binds M2 protein and interferes with the M1-M2 interaction leading to defective viral assembly ^18^.

Interestingly, M2 protein was identified as first viroporin ^30,31^. In coronaviruses several viroporins have been discovered, including SARS-CoV-1 E ^32,33^. As E proteins are highly conserved in the SARS family, we investigated a possibility that SARS-CoV-2 Envelope and cyclin D3 are potential binding partners. Our work indeed revealed cyclin D3 as a new interactor with SARS-CoV-2 E. Recently a comparative viral-human protein-protein interaction analysis for SARS-CoV-2 have been published ^34^. In their study they did not uncover cyclin D3 as an interactor with any viral protein, however, the study was conducted in HEK293T cells, cell line that we showed does not express cyclin D3. It has been shown that SARS-CoV-1 and SARS-CoV-2 proteins M, E and N are required for virion assembly that takes place in ER-Golgi intermediate compartment cisternae ^35-37^. M and E proteins seem to present an assembly core interacting with both Spike and Nucleocapsid proteins ^38,39^. Importantly, our data show that cyclin D3 associates with M as well, supporting our hypothesis that cyclin D3 impairs SARS-CoV-2 assembly and spread. Further, E and M proteins have been implicated in Spike processing and trafficking ^27^. It has been shown that Spike is retained inside cells when expressed together with E and M probably to target S to proximity of intracellular virus assembly sites. Our data show that S is retained in the cells in the presence of M and E but its trafficking towards membrane and ability to form syncytia is partially rescued when cyclin D3 is present. This supports the concept of cyclin D3 being restriction factor impairing role of M and E in SARS-CoV-2 optimal assembly.

Our work provides important insight into mechanism through which cyclin D3 limits SARS-CoV-2 infection. In the light of immune evasion from vaccination, it is important that this phenomenon was observed across different SARS-CoV-2 variants suggesting that this mechanism provides a universal target for development of antivirals. Our data suggest that cyclin D3 associates with SARS-CoV-2 E and M proteins, thereby interfering with efficient assembly. SARS-CoV-2 has therefore evolved strategies to degrade cyclin D3 that require further investigation, with the hope that it can be translated to therapeutics.

## Materials and Methods

### Reagents

#### Cell lines

All cells were maintained in Dulbecco’s modified Eagle medium (DMEM) supplemented with 10% fetal calf serum (FCS), 100 U ml^−1^ penicillin and 100 mg ml^−1^ streptomycin and regularly tested and found to be mycoplasma free. Following cells were a gift from: A549 ACE2/TMPRSS2 ^40^ Massimo Palmerini, Vero E6 ACE2/TMPRSS2 from Emma Thomson, HeLa-ACE2 from James Voss, 293T (a human embryonic kidney cell line, ATCC CRL-3216). 293T GFP11 cells and Vero-GFP10 cells for Split GFP assay were a gift from Leo James^41^.

#### Viruses

WT (lineage B, SARS-CoV-2/human/Liverpool/REMRQ0001/2020), a kind gift from Ian Goodfellow, previously isolated by Lance Turtle (University of Liverpool), David Matthews and Andrew Davidson (University of Bristol). Alpha variant (B.1.1.7; SARS-CoV-2 England/ATACCC 174/2020) was a gift from G. Towers ^42^, Delta variant (B.1.617.2) ^43^. Viral stocks were prepared by passaging once in VERO AT2 cells. Cells were infected at low MOI and incubated for 72h. Virus containing culture supernatants were clarified by centrifugation (500xg, 5min) and aliquots frozen at -80°C. Standard TCID50 assay in VERO AT2 was used to determine MOI of viral stocks.

#### Plasmids

pBOB-EF1-FastFUCCI-Puro was a gift from Kevin Brindle & Duncan Jodrell (Addgene plasmid # 86849 ; http://n2t.net/addgene:86849 ; RRID:Addgene_86849) ^29^. pCMV5 cyclin D3 HA was obtained from MRC-PPU Reagents and Services. Rc/CMV cyclin D1 HA was a gift from Philip Hinds (Addgene plasmid # 8948 ; http://n2t.net/addgene:8948 ; RRID:Addgene_8948) ^44^. pLVX-EF1alpha-SARS-CoV-2-E-2xStrep-IRES-Puro (Addgene plasmid # 141385 ; http://n2t.net/addgene:141385 ; RRID:Addgene_141385); pLVX-EF1alpha-SARS-CoV-2-M-2xStrep-IRES-Puro (Addgene plasmid # 141386 ; http://n2t.net/addgene:141386 ; RRID:Addgene_141386). pLVX-EF1alpha-SARS-CoV-2-nsp9-2xStrep-IRES-Puro (Addgene plasmid # 141375 ; http://n2t.net/addgene:141375 ; RRID:Addgene_141375); pLVX-EF1alpha-SARS-CoV-2-N-2xStrep-IRES-Puro (Addgene plasmid # 141391 ; http://n2t.net/addgene:141391 ; RRID:Addgene_141391) were a gift from Nevan Krogan ^34^. pEXN-MNCX, MLV vector encoding N-terminal double HA tag ^45^. pCAGGS_SARS-CoV-2_Spike was obtained from NIBS.

#### Antibodies

Following antibodies were used. Anti-rabbit IgG, HRP-linked Antibody (7074); Cyclin D3 Mouse mAb (DCS22, 2936); from Cell Signaling. Mouse IgG HRP Linked Whole Ab (NXA931V); from Sigma. Goat anti-Mouse IgG (H+L) Cross-Adsorbed Secondary Antibody: Alexa 488 (A-11001), Alexa 594 (A-11032), Alexa 647 (A-21236); Goat anti-Rabbit IgG (H+L) Cross-Adsorbed Secondary Antibody: Alexa 488 (A-11034), Alexa 405 (A-48254); Rabbit polyclonal SARS-CoV-2 Spike (PA1-41165); Rabbit monoclonal SARS-CoV-2 Nucleocapsid (MA5-29982) from Thermo Fisher Scientific. Mouse monoclonal Cyclin D3 (D-7, sc-6283) from Santa Cruz. Rabbit Polyclonal Cyclin A2 antibody (GTX103042); Rabbit Polyclonal Cyclin D1 antibody (N1C3, GTX108824); Rabbit Polyclonal Cyclin E1 antibody (GTX103045); Rabbit Polyclonal Cyclin B1 antibody (GTX100911); monoclonal SARS-CoV-2 Spike (GTX632604) from GeneTex. Mouse monoclonal Strep II Tag Antibody (NBP2-43735) from Novus Biologicals. Mouse monoclonal actin (ab6276) from Abcam. Strep-Tactin-HRP; MagStrep “type3” XT beads (2-4090-002) from IBA Lifesciences. Anti-HA Magnetic Beads (88836) from Thermo Fisher Scientific.

#### SDS–PAGE and immunoblots

Cells were lysed in reducing Laemmli SDS sample buffer containing PhosSTOP (Phosphatase Inhibitor Cocktail Tablets, Roche, Switzerland) at 96°C for 10 min and the proteins separated on NuPAGE^®^ Novex^®^ 4–12% Bis–Tris Gels. Subsequently, the proteins were transferred onto PVDF membranes (Millipore, Billerica, MA, USA), the membranes were quenched, and proteins detected using specific antibodies. Labelled protein bands were detected using Amersham ECL Prime Western Blotting Detection Reagent (GE Healthcare, USA) and ChemiDoc MP Imaging System (Bio-Rad) CCD camera. Protein band intensities were quantified using ChemiDoc MP Imaging System and Image Lab software (Bio-Rad, Hercules, CA, USA).

#### Immunofluorescence

Cells were fixed in 4% PFA, quenched with 50 mM NH_4_Cl and permeabilised with 0.1% Triton X-100 in PBS. After blocking in PBS/1% FCS, cells were labelled for 1 h with primary antibodies diluted in PBS/1% FCS, washed and labelled again with Alexa Fluor secondary antibodies for 1 h. Cells were washed in PBS/1% FCS and stained with DAPI in PBS for 5 min. Labelled cells were detected using ArrayScan high-content system (ThermoFisher, Waltham, MA, USA) and analysed using Harmony (PerkinElmer, Waltham, MA, USA) and ImageJ software. Infected cells have been identified by SARS-CoV-2 Nucleocapsid or Spike staining. To measure the location of cyclin D staining in cells, DAPI staining was used to demarcate the nuclear and cytoplasmic regions of interest (ROI). Harmony (PerkinElmer, Waltham, MA, USA) and ImageJ software were used to measure MFI for each protein in each region. Values are presented as a ratio of signal (nucleus/cytoplasm). Usually, at 50-200 cells have been quantified.

#### Cell cycle analysis using fluorescence ubiquitination cell cycle indicator (Fucci)

Fucci cassete was cloned from pBOB-EF1-FastFucci-Puro vector to pEXN-MNCX using BamHI/NotI restriction sites. Fucci containing lentiviral particles were produced as follows. 293Tv cells were transfected with pEXN-MNCX-Fucci, CMVi and pMD2.G. Cell supernatants containing viruses (Fucci VLP) were collected 48h post-transfection and frozen at -80°C. Cells were transduced using Fucci VLP for 18h. Cells were infected with SARS-CoV-2 variants and fixed in 4% PFA 24h post-infection. SARS-CoV-2 positive cells were identified by Nucleocapsid staining and Flow cytometry. Cell populations positive or negative for SARS-CoV-2 nucleocapsid staining were gated and Cdt1-RFP positive (G1 phase), Geminin-GFP positive (S/G2/M phase), and Cdt1-RFP/ Geminin-GFP positive (early S phase) populations were identified using flow cytometry using LSRFortessa X-20 (BD Biosciences, UK) and FlowJo software (Tree Star, OR, USA).

For immunofluorescence and high-throughput microscopy cells were transduced using Fucci VLP for 18h. Cells were infected with SARS-CoV-2 variants and fixed in 4% PFA 24h post-infection. SARS-CoV-2 positive cells were identified by Nucleocapsid or Spike staining. Cells in G1 phase were identified by Cdt1-RFP; early S, Cdt1-RFP/ Geminin-GFP; and S/G2/M, Geminin-GFP signal.

#### Knock-down

Cells were transfected with 20 pmol of siRNA Ambion Silencer Negative Control #1, predesigned Invitrogen Silencer siRNAs for cyclin D3 (siRNA ID s2523, Chr.6: 41934933 – 42048894), cyclin D1 (siRNA ID s229, Chr.11: 69641105 – 69654474), cyclin A2 (siRNA ID s2514, Chr.4: 121816444 – 121823933) using Lipofectamine RNAiMAX Transfection Reagent (Invitrogen). Medium was replaced 18h post-transfection and cells infected with Delta (MOI 0.001), Alpha and WT (MOI 0.1) variant to allow multiple rounds of infection for 4h. Cells were washed twice in PBS and incubated in new medium for 48h. Cell supernatant were collected and used to determine virus titers by standart TCID50, cells were lysed and used for western blotting to detect viral and cyclins protein expression.

#### Co-immunoprecipitation

Cells were transfected with SARS-CoV-2 genes encoding full length Spike (no tag), Strep-tagged Nucleocapsid, Envelope, Membrane, nsp9 proteins in the absence/presence of HA-cyclin D3 for 24h. Cells were lysed in Pierce IP lysis buffer (ThermoFisher, Waltham, MA, USA) supplemented with protease and phosphatase inhibitor cocktail (Pierce, Rockford, IL, USA) and 1% digitonin. Cell lysates were precleared by centrifugation. A sample of whole cell lysate was stored at this point. pre-cleared cell lysates were incubated with a-HA magnetic beads, MagStrep beads (IBA-Lifescience, Gottingen, Germany) or anti-cyclin D3 monoclonal antibody (sc-xx) bound Protein G Dynabeads for 1h at 4°C. Beads were washed 3x in IP lysis buffer and 1x in PBS. BTX elution buffer (IBA-Lifescience) was used to elute proteins from MagStrep beads. Laemmli reducing buffer was added to a-HA beads and Dynabeads and eluates from MagStrep beads and 10min at 90°C was used to elute/denaturate attached proteins. Samples were stored till further use.

#### Cell to cell fusion assay

293T GFP11 cells were transfected with WT full length Spike, and/or with WT Envelope, Membrane, cyclin D3, and empty vector (pCDNA, to ensure equal amount of transfected DNA). 24h post-transfection cells were seeded at a 1:1 ratio with Vero-GFP10 cells, final cell number 6×10e4 cells/well. Cell to cell fusion was measured 18h later and determine as a proportion of green area to total phase area using ArrayScan high-content system (ThermoFisher, Waltham, MA, USA) and analysed using ImageJ software.

## Data Availability

The authors declare that all data supporting the findings of this study are available within the article and the Supplementary Information.

## Acknowledgments

We thank Voss for HeLa ACE2; and S. Rihn for the A549-ACE2/TMPRSS2 cell, N. Matheson, and S. Marelli for full length Spike, Membrane and Envelope plasmids. This research was supported by the Cambridge NIHR BRC Cell Phenotyping Hub. In particular, we wish to thank V. Romashova for their advice and support in flow imaging. R.K.G. is supported by a Wellcome Trust Senior Fellowship in Clinical Science (WT108082AIA). We acknowledge additional support from Lister and Rosetrees Institutes.

## Author contributions

Conceived research and designed study: P.M., R.K.G. Performed experiments: P.M. Analysed data: P.M. Wrote the manuscript: P.M., R.K.G.

