## Supplementary figures for "Cell cycle independent role of cyclin D3 in host restriction of SARS-CoV-2 infection"

**Supplementary Figure 1**

**Densitometry data from cell lysates from SARS-CoV-2 infected cells**

(A,B) ImageJ was used to record densitometry of bands from Figure 1A,D. All band densities were normalised to actin and further normalised to uninfected cells (=1).

(A) VERO AT2 cell line.

(B) A549 AT2 cell line.

(C) VERO AT2 cells were infected with alpha (α) SARS-CoV-2 variant. Cells were fixed 24h post-infection and stained for viral proteins and cyclins. Arrowheads highlight un-infected cells and cyclin D/A nuclear localization. Arrowheads: Nuclear cyclin staining in uninfected cells.

**Supplementary Figure 2**

**SARS-CoV-2 infection relocalises cyclin D3 from the nucleus in HeLa Ace2 cells**

(A) HeLa cells expressing ACE2 were infected with WT and α variants and 24h later fixed and stained for SARS-CoV-2 nucleocapsid and cyclin D3.

(B) Quantification of ratio between nuclear and cytoplasm (N/C) staining of cyclin D3. Statistical analysis was performed using two-sided unpaired Student's t-tests; $^{***}p < 0.001$.

**Supplementary Figure 3**

**Effect of proteasome inhibition on SARS-CoV-2 infection**

(A) Proteasome inhibitors MG-132(1uM), Bortezomib (BZ, 1uM) were added to A549 AT2 cells 1h before infection (-1), at infection (0), 4h (4) and 24h (24) post-infection. Cells were infected with Delta SARS-CoV-2 variant and lysed 28h post-infection, viral protein Spike as measure of infection was detected by western blot.

(B) VERO AT2 cells were infected with Delta SARS-CoV-2 variant. Proteasome inhibitor Bortezomib (BZ, 1uM) was added to cells 8h later. Cells were fixed and stained 24h post-addition of inhibitor.

(C) Quantification of D3 cyclin relocalization from nucleus after infection. Uninfected (-) and SARS-CoV-2 infected cells were identified by negative/positive nucleocapsid staining. Ratio between nuclear and cytoplasm (N/C ratio) staining intensity of cyclins was measured using

ImageJ and Harmony (PerkinElmer). At least 50 cells have been counted. Ordinary two way ANOVA; ns, non-significant; $^{****}p < 0.0001$; $^{***}p < 0.001$; $^{**}p < 0.01$.

**Supplementary Figure 4**

**Depletion of cyclins by siRNA**

(A,B) D and A-cyclins were depleted using siRNA. A549 AT2 Cells were infected 18h later with Delta (Δ), Alpha (α) or wild type (WT) SARS-CoV-2 variants at MOI 0.001, 0.1, 0.1 respectively. Cells were washed 4h post-infection and new media added. Cells were collected 48h later, viral proteins and cyclins detected using western blotting.

**Supplementary Figure 5**

**Fucci cell cycle sensor**

(A) FUCCI (fluorescence ubiquitination cell cycle indicator) Cell Cycle Sensor is a two-color (red and green) indicator. Red: RFP-Cdt1 protein is expressed in G1 phase. GFP-Geminin protein is expressed in S, G2 and M phase. Both proteins are expressed in early S phase (both red and green colour).

(B) Cell cycle analysis can be performed using flow cytometry. G0/neg population of cells can not be analysed as it comprises of cells that are in G0 phase and/or were not transduced by Fucci containing lentiviral particles.

(C) Automated microscope platform and ImageJ and/or Harmony imaging software (PerkinElmer) analysis can be used to study Cell cycle changes.

**Supplementary Figure 6**

**SARS-CoV-2 infection of VERO AT2 arrests cells in S and G2/M phases**

(A-C) VERO AT2 cells were transduced with Fucci containing lentiviral particles for 18h and infected with SARS-CoV-2 WT in the absence (-) or presence of Chloroquine (CQ), Remdesivir (RVD) or infected with heat inactivated virus for additional 24h. % infected cells was determined by staining of SARS-CoV-2 nucleocapsid (NP) in infected cells using flow cytometry.

(B) Western blot for SARS-CoV-2 Spike protein as a measure of infection in cells.

(C) Analysis of cell cycle phases. n = 3; Statistical analysis was performed using two-sided unpaired Student's t-tests; ns, non-significant; $^{***}p < 0.001$; $^{**}p < 0.01$ . Bars indicate mean with SD.

(D,E) VERO AT2 cells were transduced with Fucci containing lentiviral particles for 18h and infected with SARS-CoV-2 WT at different MOI for 24h.

(D) Cells were lysed and used for Western blot.

(E) Cells and their cell cycle status were analysed using Flow cytometry. n = 3; one-way ANOVA with Dunnett's multiple comparisons test: $^{***}p < 0.001$; $^{**}p < 0.01$. Bars indicate mean with SD.

(F-G) A549 AT2 cells were transduced with Fucci containing lentiviral particles for 18h and infected with SARS-CoV-2 variants for additional 24h. Comparison of uninfected cell populations from truly uninfected cells (not exposed to virus, **uninfected**) and cells exposed to SARS-CoV-2 but uninfected (Nucleocapsid negative, **uninfected (INF)**) or **infected** (Nucleocapsid positive).

(F) Example of gating strategy for cell cycle analysis.

(G) Quantification of cell cycle arrest in early S phase after exposure of SARS-CoV-2 variants. α, alpha; Δ, delta; WT, Wuhan. n = 3; two-way ANOVA test: ns, non-significant; $^{****}p < 0.0001$; $^{***}p < 0.001$; $^{**}p < 0.01$. Bars indicate mean with SD.

**Supplementary Figure 7**

**D-cyclin knock-down arrests cell cycle in uninfected cells**

(A-C) Cyclins D1, D3 were depleted by siRNA. Cells were transduced with Fucci VSV-G pseudotype virus 18h later and flow cytometry used to identify cell cycle phases 24h later.

(A) Western blot of cell lysates from A549 AT2 cells depleted for various cyclins.

(B) Flow cytometry analysis of cell cycle comparing cyclin D1, D3 knockdown to NT (non-target siRNA) in A549 AT2 cells. n = 3; ordinary two way ANOVA with Dunnett’s multiple comparisons test: ns, non-significant; $^{****}p < 0.0001$; $^{***}p < 0.001$; $^{*}p < 0.1$. Bars indicate mean with SD.

(C) Flow cytometry analysis of cell cycle comparing cyclin D1, D3, combined D1+D3 knockdown to NT (non-target siRNA) in VERO AT2 cells. n = 3; ordinary two way ANOVA with Dunnett’s multiple comparisons test: ns, non-significant; $^{**}p < 0.01$; $^{*}p < 0.1$. Bars indicate mean with SD.

**Supplementary Figure 8**

**SARS-CoV-2 mediated depletion of D-cyclins is cell cycle arrest independent**

(A-D) A549 AT2 cells were depleted for D and A2 cyclins and 18h later infected with Alpha variant SARS-CoV-2 for 24h. Cells were fixed, stained for SARS-CoV-2 nucleocapsid and analysed for infection and Fucci cell cycle sensor.

(A) Percentage of infected cells in cells depleted for cyclins. n=2; Ordinary two way ANOVA with Sidak's multiple comparisons test: ns, non-significant; $^{****}p < 0.0001$; $^{***}p < 0.001$; $^{*}p < 0.1$. Bars indicate mean with SD.

(B) Flow cytometry analysis of early S cell cycle phase comparing cyclin D1, D3, and A2 knockdown to NT (non-target siRNA) in two independent experiments in duplicates.

Statistical analysis was performed using two-sided unpaired Student's t-tests; ns, non-significant; $^{***}p < 0.001$; $^{**}p < 0.01$; $^{*}p < 0.1$. Bars indicate mean with SD.

(C,D) Comparison of uninfected cell populations from truly uninfected cells (not exposed to virus, **uninfected**) and cells exposed to SARS-CoV-2 but uninfected (Nucleocapsid negative, **uninfected (INF)**). A549 AT2 cells were transduced with Fucci containing lentiviral particles for 18h and infected with (C) Alpha and (D) Delta SARS-CoV-2 variants for additional 24h.

(E-H) VERO AT2 cells were transduced with VSV-G pseudotyped Fucci containing lentiviral particles and 18h later infected with Alpha SARS-CoV-2. Cells were fixed and stained for SARS-CoV-2 nucleocapsid, D-cyclins and analysed for infection and Fucci cell cycle sensor 24h later. Statistical analysis was performed using two-sided unpaired Student's t-tests; ns, non-significant. Bars indicate mean with SD.

(E) VERO AT2 cells infected with Alpha variant. Flow cytometry analysis of cell cycle comparing cyclin D1, D3, combined D1+D3 knockdown to NT (non-target siRNA) in uninfected and SARS-CoV-2 infected cells. Plot is an example of 3 independent experiments, in duplicates. Statistical analysis was performed using two-sided unpaired Student's t-test: ns, non-significant; $^{****}p < 0.0001$. Bars indicate mean with SD.

(F) VERO AT2 cells infected with Delta variant. Flow cytometry analysis of S/G2/M cell cycle phase comparing cyclin D3 knockdown to NT (non-target siRNA). n = 3. Statistical analysis was performed using two-sided unpaired Student's t-test: ns, non-significant; $^{***}p < 0.001$; $^{**}p < 0.01$; $^{*}p < 0.1$. Bars indicate mean with SD.

(G) Example of acquisition using automated microscopic platform. Cells are identified for infection, cell cycle (Red/arrow=G1phase; Green/arrowhead=S/G2/M; Red+Green/arrowhead=early S) and expression of cyclin D3.

(H) Quantification of cyclin D3 re-localization from nucleus to cytoplasm and correlation with cell cycle phases using ImageJ and Harmony (PerkinElmer). At least 50-200 cells were analysed in each condition. Statistical analysis was performed using two-sided unpaired Student's t-test: ****$p < 0.0001$; ***$p < 0.001$.

**Supplementary Figure 9**

**Cyclin D3 associates with E and M proteins**

(A) Western blot of cell lysates from cell lines used in this study, detecting endogenous expression of cyclin D1, D3.

(B) 293T cells were contransfected with HA-cyclin D3 and SARS-CoV-2 Spike, or Strep-tagged E, M, N or nsp9. Whole Cell lysates show expression levels of protein input into immunoprecipitation.

(C-E) Immunoprecipitation was performed using (B) mouse anti-cyclin D3 antibody, (C) anti-HA antibody, (D) anti-Strep beads. The immunoprecipitates were blotted and stained with anti-Strep, anti-HA, and anti-Spike antibodies. *non-specific band.

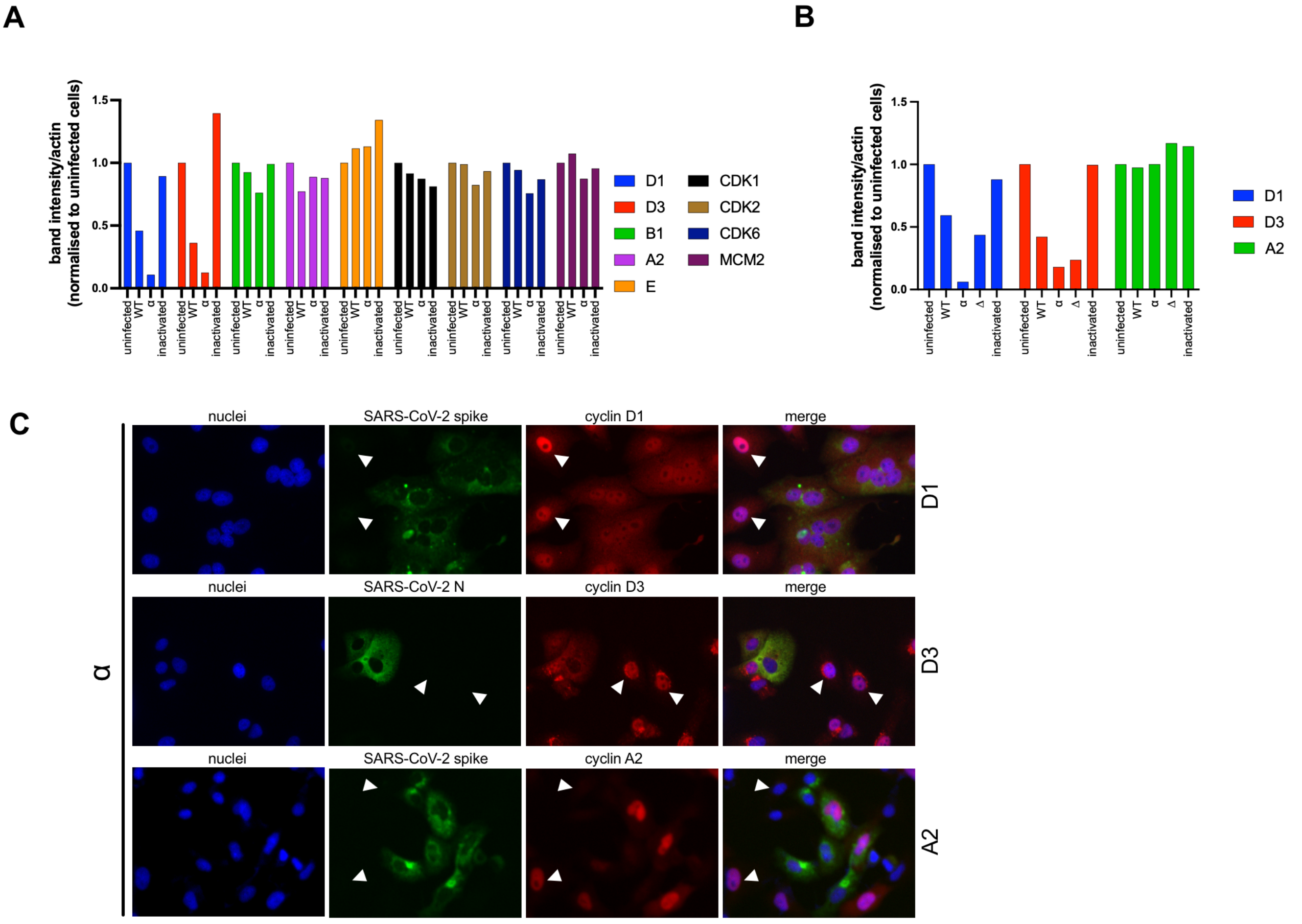


Supplementary Figure 1

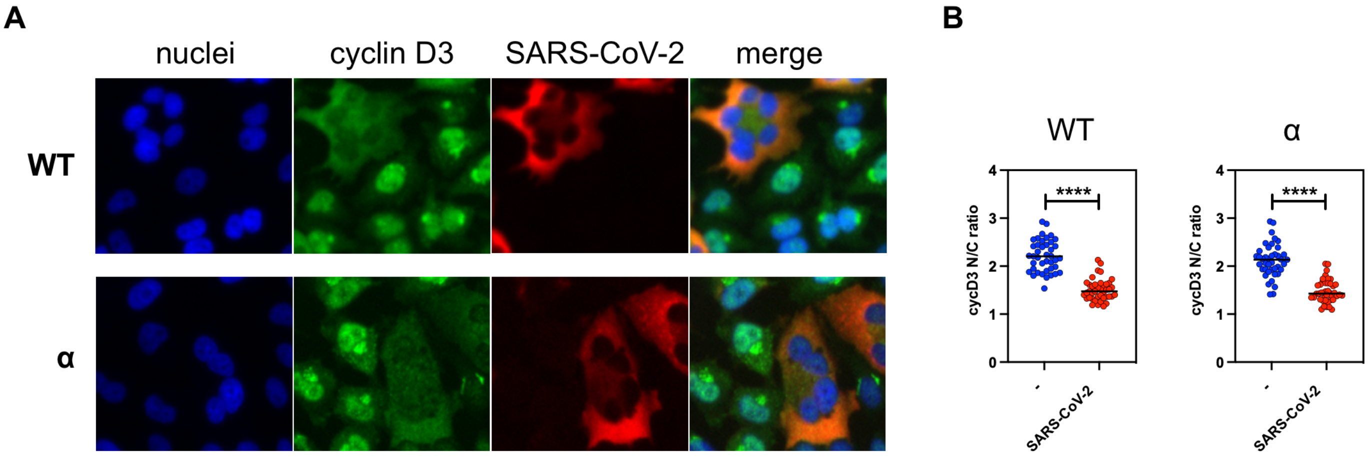


Supplementary Figure 2

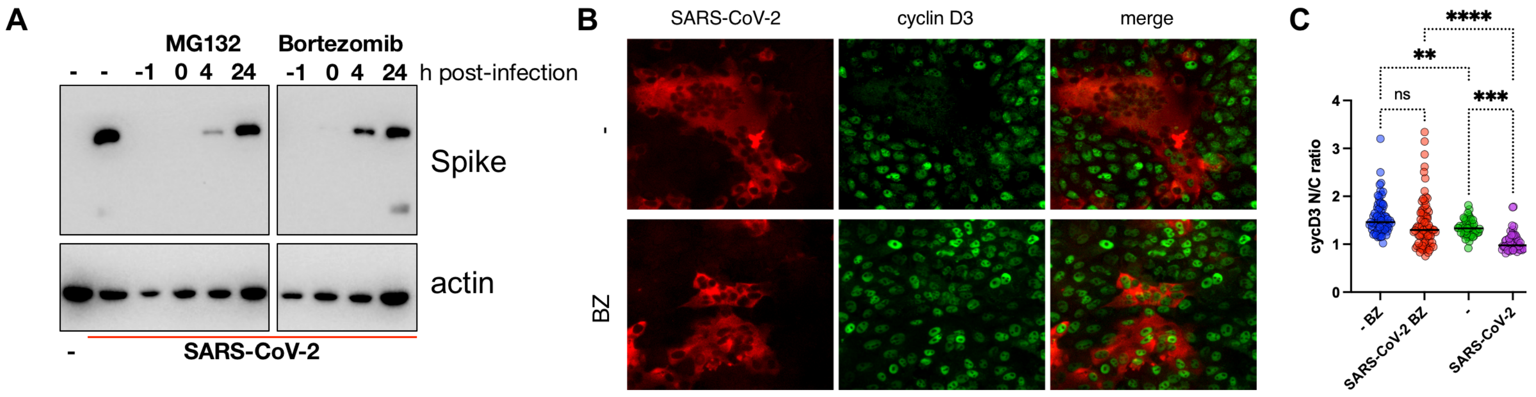


Supplementary Figure 3

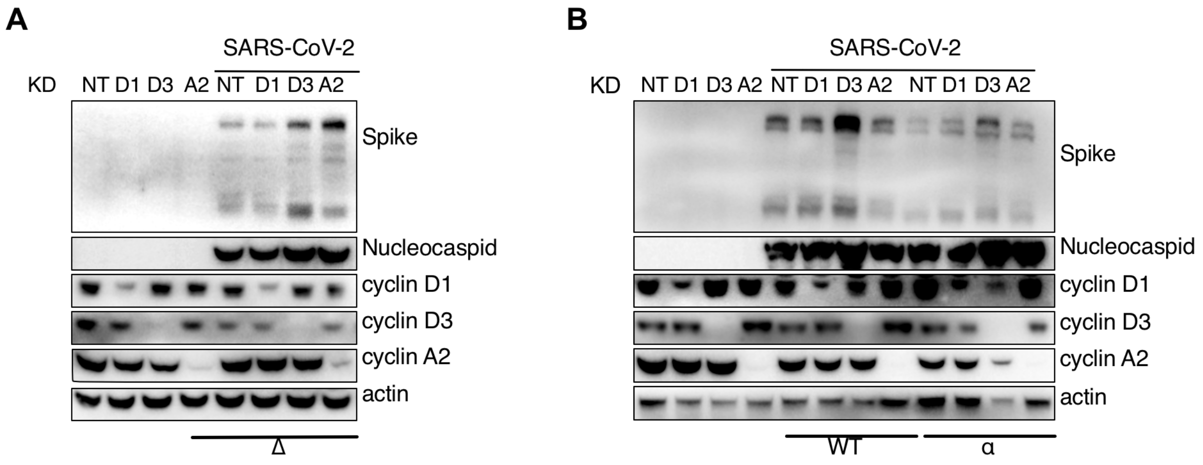


Supplementary Figure 4

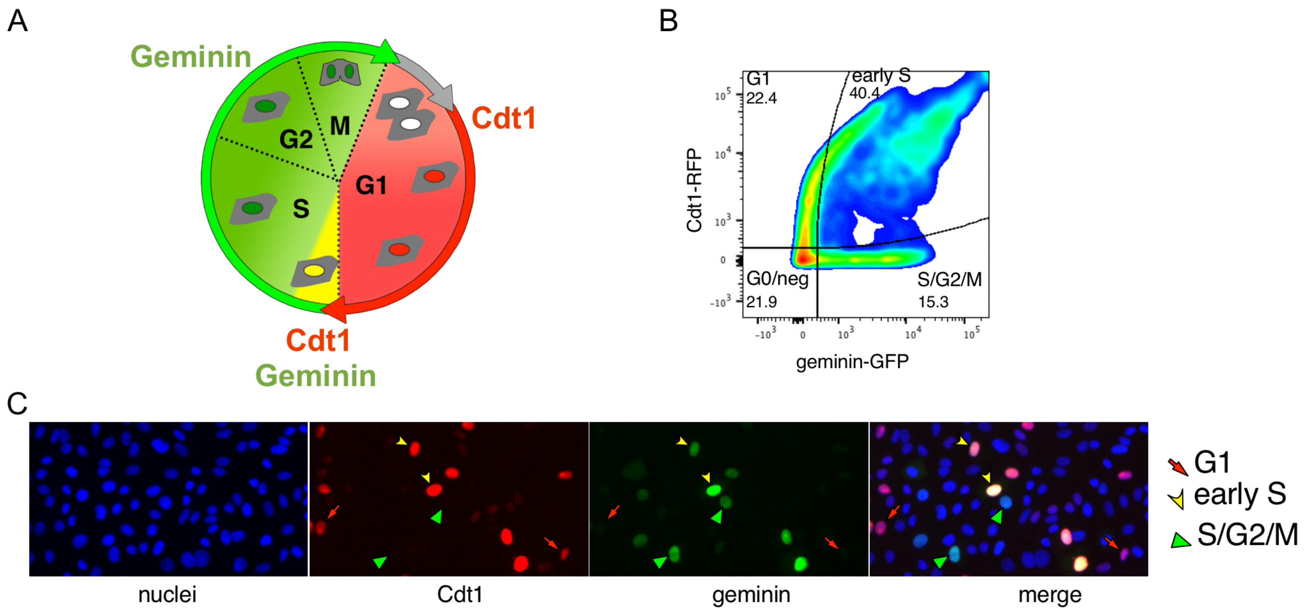


Supplementary Figure 5

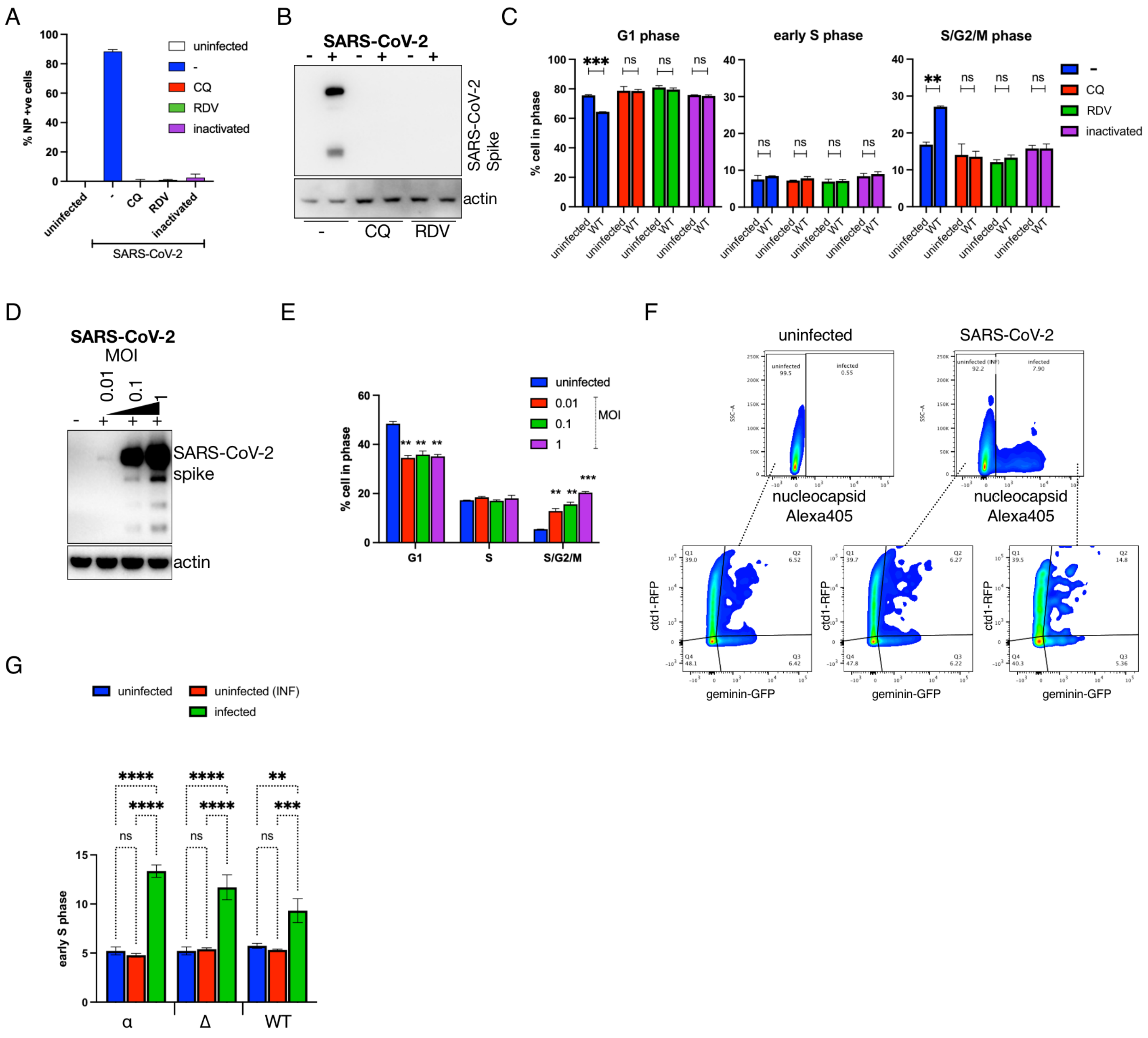


Supplementary Figure 6

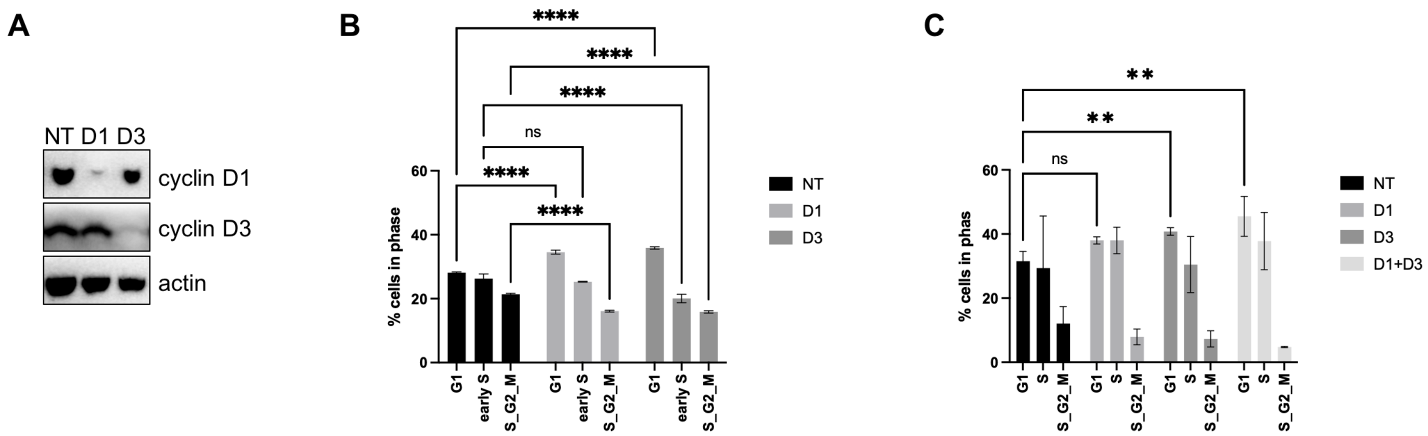


Supplementary Figure 7

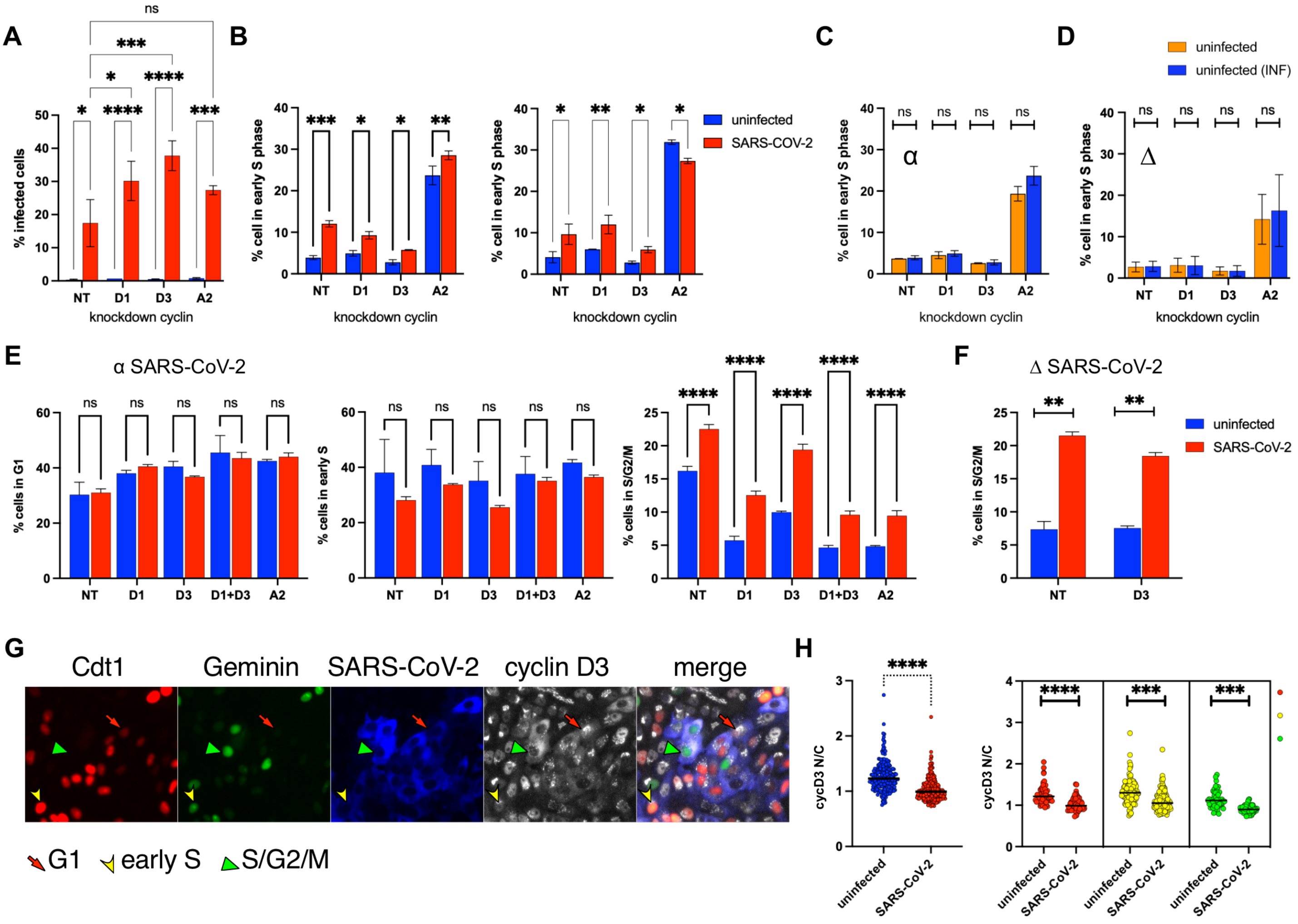


Supplementary Figure 8

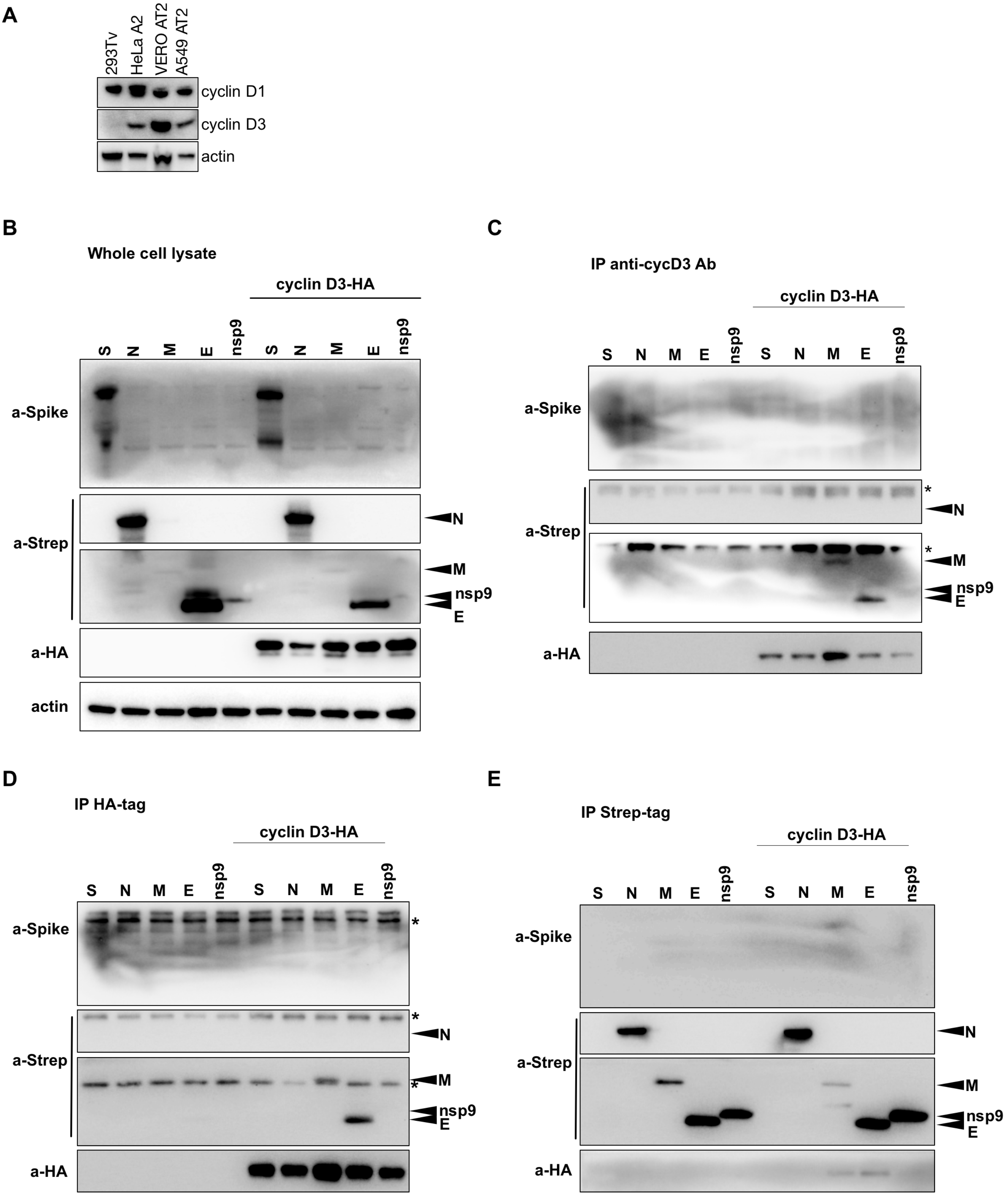


Supplementary Figure 9
